# Effect of pretreatment with different MMP inhibitors on shear bond strength of composite to primary teeth dentin after six months of aging

**DOI:** 10.1101/866129

**Authors:** Zahra Parsaie, Najmeh Mohammadi, Maryam Firouzmandi

## Abstract

**Background:** loss of hybrid layer integrity compromises the resin-dentin bond stability. Subsequent release and activation of matrix metalloproteinase enzymes during dental restorative procedures or caries development are contributing factors to dentin-adhesive bond failure.

**Aim:** To investigate the effect of pre-treatment with different MMP inhibitors on the shear bond strength (SBS) of an etch-and-rinse adhesive (Adper Single Bond 2) to primary teeth dentin after six months of aging.

**Methods:** Sixty extracted anterior primary teeth, due to orthodontic reasons, were selected. A dentin block (6.0 mm × 6.0 mm × 2.0 mm) was obtained from each tooth. All the dentin blocks (n = 60) were pretreated for 60 s with either 0.01 M phosphate buffered saline (PBS, pH 7.2) in the control group, 2%: Chlorhexidine (CHX) solution, 2% Doxycycline (DO)solution and EDTA 17% after etching and before applying etch-and-rinse adhesive system (Adper Single Bond 2). Then composite was applied in clear teflon cylinders. The specimens were stored in artificial saliva for 6 months at 37°C and then the SBS values were determined with a universal testing machine. Representative samples were selected for visualization of the failure mode under stereomicroscope and SEM.

**Results:** No statistically significant difference observed between the SBS of the EDTA group, DO group and the control group (P value>0.05). CHX showed significantly higher SBS values compared to the other groups of the study after six months(P value=0.011).

**Conclusion:** Chlorhexidine significantly preserved the SBS of composite resin to deciduous dentin using etch and rinse adhesive Adper Single Bond 2 after 6 months of aging in artificial saliva.

## Introduction

Long-term studies have shown a decrease in the bond strength of resin-bonded dentin over time [1, 2]. Degradation of the hybrid layer at the dentin-adhesive interface counts as the primary reason for this phenomenon [3]. Indeed, loss of hybrid layer integrity compromises the resin-dentin bond stability [1].

Apart from extrinsic factors, such as water or oral fluid sorption and polymer swelling, intrinsic host-derived matrix metalloproteinases (MMPs) are also involved in the breakdown of the hybrid layer [4].

MMPs are a class of zinc and calcium dependent endopeptidases which are trapped within the mineralized dentin matrix during the tooth development process. The subsequent release and activation of these enzymes during dental restorative procedures or caries development are considered as contributing factors to dentin-adhesive bond failure [5].

Simplified etch-and-rinse adhesives as well as the less destructive versions of self-etching adhesives have been verified with the capability to release and activate endogenous MMPs during the dentin bonding procedure [4]. Moreover, etch-and-rinse adhesives might lead to the formation of incompletely infiltrated zones and denuded collagen fibrils along the bottom of the hybrid layer as a result of the decreasing gradient of resin monomer diffusion within the acid-etched dentin [6]. Dentin matrix metalloproteinases can devastate these unprotected collagen fibrils [7]. Thus, from a clinical standpoint, MMP inhibitors, such as chlorhexidine (CHX) can play an imperative role in the longevity of the resin bond to dentin [3].

CHX acts by cation chelation, sequestrating calcium and zinc ions essential for the activation of the MMPs and inhibiting collagenase and gelatinase activity of dentin matrix [8]. Recent *in vivo* and *in vitro* studies have demonstrated that application of chlorhexidine has broad-spectrum MMP-inhibitory effects and significantly improves the integrity of the hybrid layer created by a simplified etch-and-rinse adhesive [4].

In addition, antibiotics such as Tetracyclines (TCs) and their semisynthetic forms namely doxycycline (DO) are commonly used in the treatment of periodontitis and have the ability to inhibit the collagen degradation activity of MMPs. Following the etching procedure, pre-treatment of the dentin surface with an aqueous solution of 2% doxycycline can inhibit collagenase and gelatinase enzymes [5].[9]

Based on the findings of a recent research, applying aqueous solutions of semisynthetic TCs (minocycline and doxycycline) as pre-treatment following the acid-etching procedure results in an improvement of immediate bonding performance [10].

Another extrinsic agent with MMP inhibitory capacity is ethylenediaminetetraacetic acid (EDTA). The application of EDTA as a chelating compound has been shown to inactivate the endogenous MMP action in human dentin [5]. Singh *et al*., in their study, showed that EDTA has a MMP inhibitory effect which might help to enhance the durability of resin-dentin bond [11].

Beneficial effects of dentin surface pre-treatment with MMP inhibitors would possibly become clear over the course of time, as long as the dentin bond strength is not immediately impaired [12].

However, to date, the efficacy of these MMP inhibitors to prevent loss of dentin bond strength over time has not been determined in primary dentition. Thus, the aim of the present study was to investigate the effect of pre-treatment with different MMP inhibitors on the shear bond strength (SBS) of an etch-and-rinse adhesive (Adper Single Bond 2) to primary teeth dentin after six months of aging along with microscopic evaluation of the bond failure mode.

## Materials and Methods

### Initial specimen preparation

The local ethics committee of Shiraz University of Medical Sciences approved the study protocol (Grant#20969). Then sixty primary anterior teeth were provided from teeth extracted due to orthodontic reasons. Written informed consent was obtained from parents or guardians at the time of tooth extraction and the they were informed about using the samples for research purposes. The selected teeth were inspected visually and confirmed as those free of discoloration, carious lesions, and any defect. The enamel was removed to create a flat dentinal surface and then the root was cut at the cementum-enamel junction. A dentin block (6.0 mm × 6.0 mm × 2.0 mm) was obtained from each tooth. Lack of enamel residue was confirmed using a stereomicroscope (BS-3060C, BestScope, China). Next, the dentin blocks were polished with a #600-grit wet silicon carbide abrasive paper. Then, the samples were rinsed thoroughly with water.

### Etching and bonding procedures and treatment groups

For the acid conditioning procedure, 37% phosphoric-acid gel (Scotchbond Etchant, 3M ESPE) was used for 20 s and further washed with water for 15 s. Dentin was then pretreated with three different MMP inhibitors. All the dentin blocks (n = 60) were pretreated for 60 s with either 0.01 M phosphate buffered saline (PBS, pH 7.2) in the control group, 2% CHX solution, 2% Doxycycline solution and EDTA 17% after etching and before applying etch-and-rinse adhesive system (Adper Single Bond 2).

The four groups inspected in this study were as follows:

group I- control group (n=15): 0.01M phosphate buffered saline of pH 7.2 was used. Dentinal surfaces were dried with the aid of an absorbent and air stream. Then, the resin adhesive layer was applied.

Group II-chlorhexidine (n=15): 2% chlorhexidinegluconate solution was applied for 60 s on the dentin blocks. The application of CHX was performed using a micro-brush. Dentinal surfaces were dried with the use of an absorbent and air stream. Then, the resin adhesive layer was applied.

Group III- Doxycycline (n=15):2% doxycycline solution was applied for 60s on the dentin blocks. The application of DO was conducted by means of a micro-brush. Dentinal surfaces were dried with the use of an absorbent and air stream. Then, the resin adhesive layer was applied.

Group IV: EDTA (15 samples):EDTA17% was applied for 60 s on the dentin blocks. The application of EDTA was perfomed by means of a micro-brush. Dentinal surfaces absorbent and air stream. Then, the resin adhesive layer was applied.

Following etching, pre-treatment, and adhesive application in accordance with the manufacturer instructions (table 1), composite was applied. In all four groups, a clear plastic mold (Tygon tubes, ET, Shandong China) measuring 2 mm in diameter and 2 mm in length was secured to the lapped tooth surface and served as a mold into which the composite (Filtek 3M,USA) was inserted. The composite was cured for 20 s according to the manufacturer instructions from three different sides (one from the top and two from the sides). The specimens were stored in artificial saliva for 6 months at 37°C and then the shear bond strength values were determined with a universal testing machine (Zwick/Roll Z020, Zwick GmbH & Co, Germany). The test was performed by securing the specimens in a mounting jig and a sharp straight-edge chisel attached to the cross-head was used to apply a shearing force of 0.5 mm /min until failure.

**Table 1:** Acid etching and adhesive application procedures.

|  | Acid etching | Adhesive application |
| --- | --- | --- |
| Material | Acid etch(Scotchbond Etchant, 3M ESPE) | Adper Single Bond 2(3M ESPE,USA) |
| Composition | Acid phosphoric 37% | Bis-GMA,HEMA,water, dimethacrylates,ethanol, photoinitiator system, methacrylate functional copolymer of polyacrylic and polyitaconic acids, 10% by |
|  |  | weight of 5 nm-diameter spherical silica nanoparticles |
| Application Technique | 1. Apply etchant for 20s<br>2. Rinse for 15s<br>3. Gently air-dry (10s at 20cm) | 1. Apply one coat of adhesive with gentle agitation<br>2. Gently air-dry (10s at 20cm)<br>3. Apply second coat of adhesive.<br>4. Gently air-dry (10s at 20cm)<br>5. Light curing for 20s |

### Mode of fracture failure

Modes of failure were examined under Stereomicroscope (BS-3060C, BestScope, China) for all specimens. Failures were categorized as one of the following:

Type I: adhesive failure in the tooth-composite interface

Type II: cohesive failure in the composite or dentin structure

Type III: mixed adhesive and cohesive failure

### Preparation for visualization using field-emission scanning microscope (SEM)

Two cut sections of sheared dentinal surfaces of each group of the study were examined using magnifications of up to 1130 X for analysis of sheared dentinal surfaces, with emphasis on areas of adhesive failure or areas of cohesive failure. The specimens were mounted on aluminum stubs with conductive silver liquid, gold sputter-coated and examined under a field-emission SEM (TE-SCAN, VEGA3, USA) for verification of type of failure.

### Statistical Analysis

SPSS version 20 (SPSS Inc., IL, USA) was used to assess the collected data. Data were analyzed using Duncan post hoc test. P values <0.05 were considered to be statistically significant.

## Results

The mean SBS values and their respective standard deviations for different experimental groups are presented in Table 2.

**Table 2:** mean SBS values ±standard deviations and failure modes for different experimental groups.

| Group | | Number | Minimum | Maximum | Mean±SD | P.value | Failure mode | | |
| --- | --- | --- | --- | --- | --- | --- | --- | --- | --- |
|  |  |  |  |  |  |  | I | II | III |
| I | Control | 15 | 3.23 | 9.45 | 6.44±2.15<sup>a</sup> | 0.011 | 10 | 0 | 5 |
| II | EDTA | 15 | 3.54 | 11.50 | 7.45±2.87<sup>a</sup> | | 4 | 6 | 5 |
| III | DO | 15 | 4.65 | 12 | 8.37±2.73<sup>ab</sup> | | 4 | 3 | 8 |
| IV | CHX | 15 | 5.47 | 14.60 | 11.18±2.86<sup>b</sup> | | 3 | 3 | 9 |

mean values with at least a common letter were not statistically different (Duncan's Post –hoc test)

One-way ANOVA revealed a statistically significant difference between the SBS of different experimental groups (P =0.011). Thus, Duncan multiple comparison test was used for Post –hoc analysis. According to the findings, there were no statistically significant differences between the mean SBS value of the control, EDTA, and DO groups (P value>0.05). CHX showed higher SBS values compared to the other study groups, with the difference being significant for EDTA and the control group.

### Microscopic analysis

The prevalence of the failure modes confirmed by Stereomicroscope are shown in Table 2. The most commonly occurring failure modes were mixed failures. Figure 1 shows the images related to the Stereomicroscope evaluation for all groups of the study. The results of SEM analysis were also consistent with the mentioned data analysis. Figure 2 illustrates representative images from visualization of cut sections of sheared dentinal surfaces by SEM. In images I (CHX group) and II (control group), which are related to the groups with highest and lowest SBS values, mixed failure and adhesive respectively are evident.

**Figure 1:**
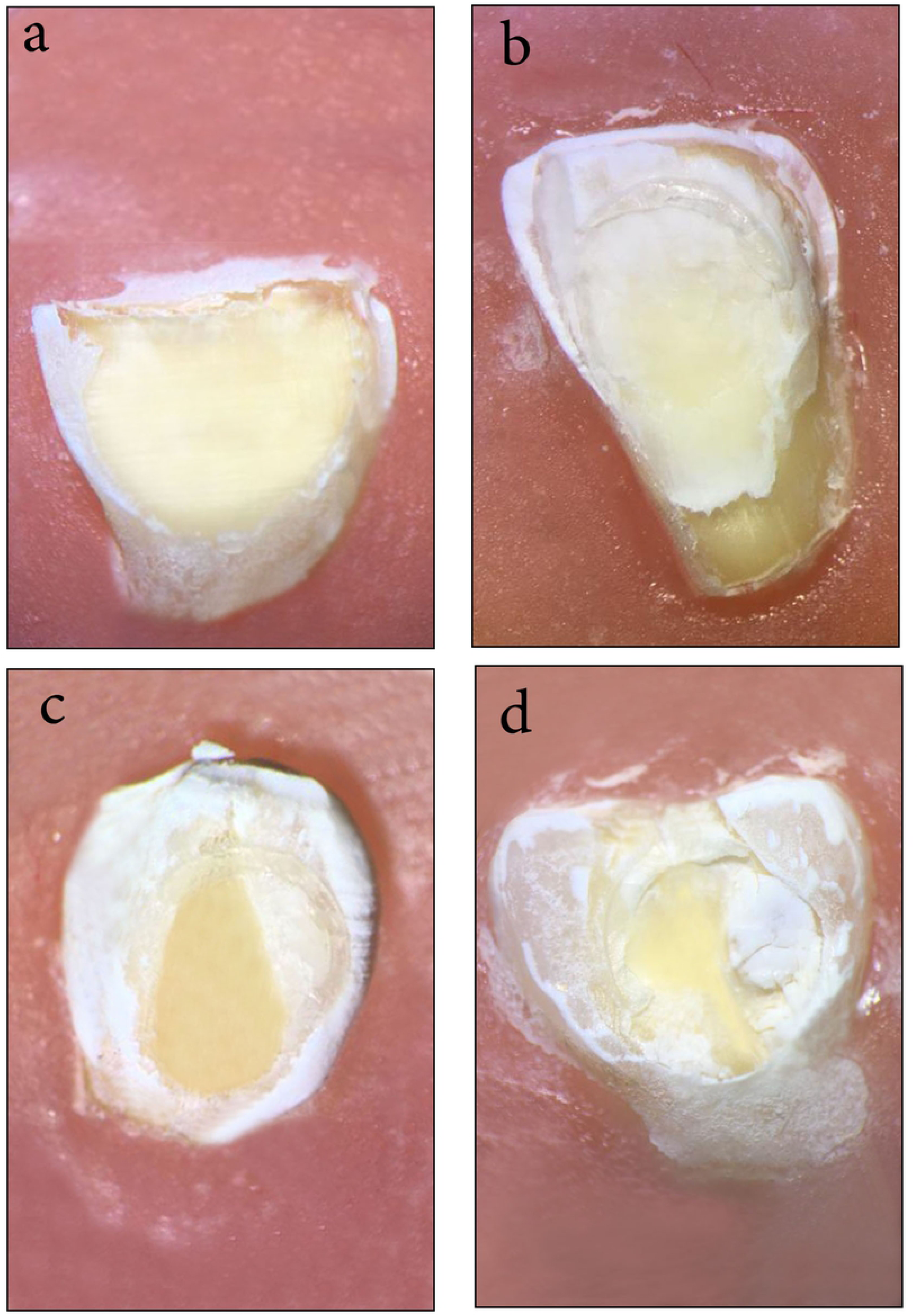
Images related to the Stereomicroscope evaluation for all groups of the study.a:control group,b:EDTA group,C:DO group,D:CHX group.

**Figure 2:**
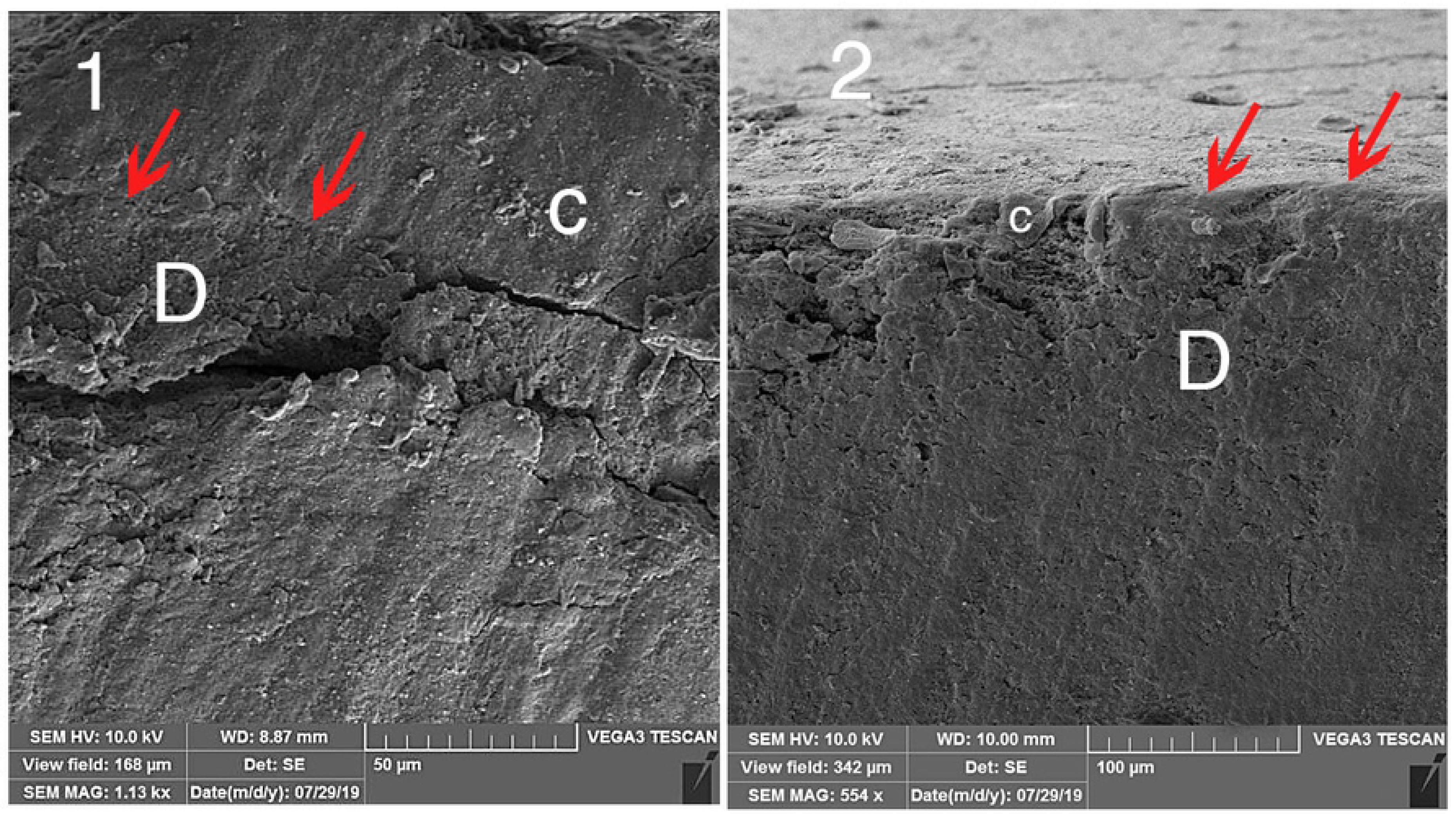
Representative images from visualization of cut sections of sheared dentinal surfaces by SEM. Images I (CHX group) and II (control group).C:Composite,D:Dentin

## Discussion

Numerous factors could affect the bond strength of adhesive agents. Mechanical stresses induced by chewing forces, temperature and pH fluctuations of the oral cavity, water sorption, resin shrinkage, and enzymatic action of MMPs are amid factors that can influence the bond integrity to diverse extents [13]. Despite the simplification of adhesive systems in recent years, etch- and -rinse adhesives are still considered as the gold standard agents regarding the durability and strength of the bond [10].The acidic action during the bonding procedure of these adhesives exposes fibrils of the organic matrix.

In order to form the hybrid layer, these fibrils must be completely infiltrated with the adhesive resin. However, a layer of denuded collagen might be left at the base of the hybrid layer as the result of insufficient infiltration of the resin monomers that are susceptible to degradation by MMP host-derived enzymes activated through acid contact and water uptake [14].[14]

Pretreatment with MMP inhibitors is considered as a valid alternative in order to prolong the resin-dentin bond stability by overcoming this self-degradation process [11].

Our study was carried out on deciduous teeth which are less mineralized than permanent teeth. They contain higher amounts of organic materials. Consequently, more degradation of the hybrid layer is probable for these teeth over time and application of MMP inhibitors seems more beneficial in these teeth [14]

The current study showed improved dentin bond strength with post-etching chlorhexidine application following 6 months of aging.Other researches have also showed that the application of 2% chlorhexidine digluconate solution before composite bonding successfully preserved the bond strength up to 6 months when etch-and-rinse adhesive systems were used [15-17]. Manfro *et al.* and Breschi *et al*. have also reported bond strengths in the CHX-treated specimens to be significantly greater than the non-treated specimens after 12 months of aging [18, 19]. The results of the current study corroborated with these findings.

The preserved bond interface associated with the use of CHX can be explained by the inhibitory ability of CHX on the matrix metalloproteinases (MMPs) found in etched dentin [7] like the results of the study by Carrilho *et al and* Zheng *et al [20, 21]*.

CHX maintains a strong affinity to the dental tissue through binding to the phosphate groups of mineralized dentin crystallites and negative carboxyl groups of the collagen matrix. Hence, long-term effectiveness of CHX would be explained by its high substantivity, regardless of its concentration. After oversaturating proteases’ binding sites, it can remain bound to collagen fibrils for a later release if still available in higher concentrations [8].

Furthermore, chlorhexidine water-based solutions can act as a rehydrating agent for dried mineralized dentin by preserving the necessary humidity needed to maintain the collagen network in an expanded condition [3].

Clinical studies like the study by Ricci *et al*. have also confirmed the results of laboratory researches regarding this MMP inhibitor [22]. An *in vivo* study on primary teeth showed extensive nanoleakage and degradation of hybrid layer even in teeth with clinically intact cavosurface margins after only 6 months of clinical service. Conversely, when CHX was applied on etched-dentin, the hybrid layer deterioration was significantly reduced [23].

Contradictory results have been reported in few studies on primary dentition. Abdelmegid reported that 2% chlorhexidine gluconate applied for 20 or 40 seconds before acid etching or conditioning did not significantly affect SBS to caries affected dentin of primary teeth[24]. Bahrololoomi *et al.* also reported that pretreatment with chlorhexidine and sodium hypochlorite increased microleakage in composite restorations in primary teeth [25]. The difference observed could be explained by implementation of different materials and methods along with pretreatment of caries affected dentin surface.

EDTA is another commonly used MMP inhibitor. The study by Thompson *et al.* asserted that 17% EDTA significantly inhibits endogenous MMP activity of human dentin within 1–2 minutes [26].

However, in the current study, the application of EDTA did not show significant differences in SBS in comparison with the control group. EDTA is water soluble; hence, it might be rinsed off the EDTA-treated dentin. This might not be able to sustain MMP inhibition for long durations such as the 6-month period implemented in this study [8].

Based on our findings, the SBS of composite to primary teeth dentin were comparable for the CHX and DOX groups after 6 months as it was reported in the study by Li and his colleagues [10]. Loguerico *et al.* also showed that the use of 2% minocycline as a pretreatment method of acid-etched dentin retarded the degradation of resin-dentin interface over a 24-month period similar to that of 2% chlorexidine digluconate [27].

However, in some other studies, doxycycline negatively affected the bonding of the etch-and-rinse adhesive [28, 29]. The increased depth of demineralization which was not compatible with the depth of resin monomer infiltration in the study by Elkassas et al. [28] and phase separation observed for doxcycycline after mixing with bonding agents in the study by Stanislawczuk et al. [29], resulted in lower shear bond strength values observed.

Evaluating the sheared surfaces by Stereomicroscope and SEM in the present study revealed that the failure modes were mostly adhesive for the control group with an increased percentage of mixed failure for MMP inhibitors. The increased percentage of mixed failures for MMP inhibitors could be attributed to the increased shear bond strength values.

Although the performance of CHX was superior in our study, it should be mentioned that the molecules of this inhibitor are water soluble and may be gradually leached out from the adhesive interface, especially when it comes in contact with an external environment (through marginal gaps, for instance). Therefore, more long-term laboratory and clinical studies (more than 6 months) are suggested for more certain results.

## Conclusion

MMP inhibitors (chlorhexidine, EDTA and doxycycline) preserved the SBS of composite resin to deciduous dentin using etch and rinse adhesive Adper Single Bond 2 after 6 months of aging in artificial saliva. However, the results were only statistically significant in the CHX group. Further *in vivo* trials should be carried out to confirm the role of CHX in bond durability.

## Acknowledgments

The authors thank the Vice-Chancellery of Shiraz University of Medical Sciences for supporting this research (Grant#19805).The authors also thank Dr. Vossoughi from the Center for Research Improvement of the School of Dentistry for statistical analysis and Farzaneh Rasouli for improving the use of English in the manuscript.

